# Dimerization of natively folded molecules drives misfolding and aggregation in Cys-depleted variants of cataract-associated human lens γD-crystallin

**DOI:** 10.64898/2025.12.29.696901

**Authors:** Yu Pu, Vanie Seecharan, Loy Hashimoto, Aslam Uddin, Eugene Serebryany

## Abstract

Cataract, the leading cause of blindness worldwide, results from age-related misfolding and aggregation of long-lived crystallin proteins in the eye lens. The cytoplasm of fiber cells in the lens core becomes increasingly oxidizing with age, allowing non-native disulfides to drive light-scattering aggregation of γ-crystallins. Despite this vulnerability to non-native disulfides, and despite lacking any native-state disulfides, γ-crystallins are unexpectedly Cys-rich. To understand this paradox, we investigated how replacing all four Cys residues in the aggregation-prone N-terminal domain of γD-crystallin affects its stability and aggregation. Cys removal precludes the disulfide-driven aggregation pathway we reported previously. Here, we characterize two full-length human γD-crystallin variants: C18S/C32S/C41S/C78S (“NCS”) and C18T/C32A/C41A/C78A (“NCA/T”). Thermodynamic and kinetic stability measurements indicate the N-terminal domain was greatly destabilized in both variants relative to WT, with NCS more destabilized than NCA/T. Upon mild heating or partial denaturation, both variants formed light-scattering aggregates, which were amorphous by transmission electron microscopy. Surprisingly, the aggregation proceeded exclusively from a dimer of natively folded molecules held together by a C-terminal disulfide bridge. These dimers form readily even in the WT protein, and evidence of them has been found in the lens. Aggregation was strongly suppressed by the lens’s native chemical chaperone, *myo*-inositol. The aggregation rate depended linearly on protein concentration, indicating that the rate limiting step was a transformation of the natively-folded to misfolded molecules within the dimer. We propose that many age-related chemical modifications could destabilize the native fold of human γD-crystallin, favor misfolding within disulfide-bridged dimers, and thereby cause aggregation.

## Introduction

Age-onset cataract is the leading cause of blindness worldwide, with no approved non-surgical treatment (1). It is primarily driven by age-related aggregation of crystallin proteins in the eye lens (2-5). Long-lived α-, β-, and γ-crystallins comprise ∼90% of total lens protein and serve to maintain transparency and refractive function for a lifetime (6-8). Since they are never replaced, these protein molecules continuously accumulate damage including deamidation, truncation, and oxidation, throughout life (9-16). In humans, γ-crystallins have only been found in the eye lens so far, although non-lenticular expression, especially of γS-crystallin, has been reported in rodents (17-19). Cataract occurs when light-scattering crystallin aggregates become comparable in size to the wavelengths of visible light (20-23). This transition correlates with the cytoplasmic redox potentials of the lens core fiber cells becoming increasingly oxidizing (24-26), and the aggregates themselves contain many non-native disulfide bonds (4, 27, 28). We and others have previously reported that formation of non-native disulfides traps γ-crystallins in misfolded conformations *in vitro* (29-31) and likely also *in vivo* (32-34). The γ-crystallins therefore present a paradox. They are Cys-rich yet contain no disulfides natively, whereas formation of non-native disulfides drives light-scattering aggregation and cataract (35). Why, then, have the γ-crystallins not evolved to become Cys-depleted, like the α-crystallins?

Redox homeostasis played an important role in the evolution of lens longevity; for example, we have reported bioinformatic evidence that disulfide avoidance shaped the evolution of γ-crystallin surfaces (35). In addition to being associated with age-onset cataracts, high oxidative stress, such as from heavy metal exposure, can cause cataracts at a younger age (36-38).

We and others have also reported oxidoreductase activity of surface Cys residues in γ-crystallins (30, 31, 39, 40). However, it is oxidation of natively buried Cys residues within the N-terminal domain (NTD) of human γD-crystallin (HγD) that drives light-scattering aggregation (29, 32, 41). Human γD-crystallin variants with a destabilized NTD have been reported to escape the aggregation-suppression function of the α-crystallin chaperones (42, 43). However, the misfolding and aggregation driven by formation of non-native disulfides within the NTD of human γD-crystallin can be delayed and mitigated by the lens’s native chemical chaperone, *myo-*inositol (44).

Here we investigate the effects of replacing all four buried NTD Cys residues in HγD (**Figure 1**) with non-disulfide forming residues. We characterize two full-length constructs: C18S/C32S/C41S/C78S (“NCS”) and C18T/C32A/C41A/C78A (“NCA/T”). NCS replaces all four NTD Cys residues with Ser, which is generally considered to be the most chemically and genetically conservative substitution. (NCS differs from WT solely by the replacement of four S atoms with O atoms.) It may also serve as a mimic of the mildest form of Cys oxidation: conversion of the thiol group (SH) to sulfenic acid (S-OH). However, Ser is more flexible (45) and more hydrophilic (46, 47) than Cys, and it risks being a frustrated hydrogen bond donor when placed in the hydrophobic protein core, so it could generate denaturation-from-within. Therefore, we tested an alternative construct, NCA/T. In this construct, we replaced the natively buried Cys32, 41, and 78 with Ala to better preserve the hydrophobic character of these sites. However, Cys18 is close enough to the surface that we chose to substitute it with Thr rather than Ala, to avoid generating any new hydrophobic surface patch and to preserve the possibility of a favorable protein-solvent interaction.

**Figure 1:**
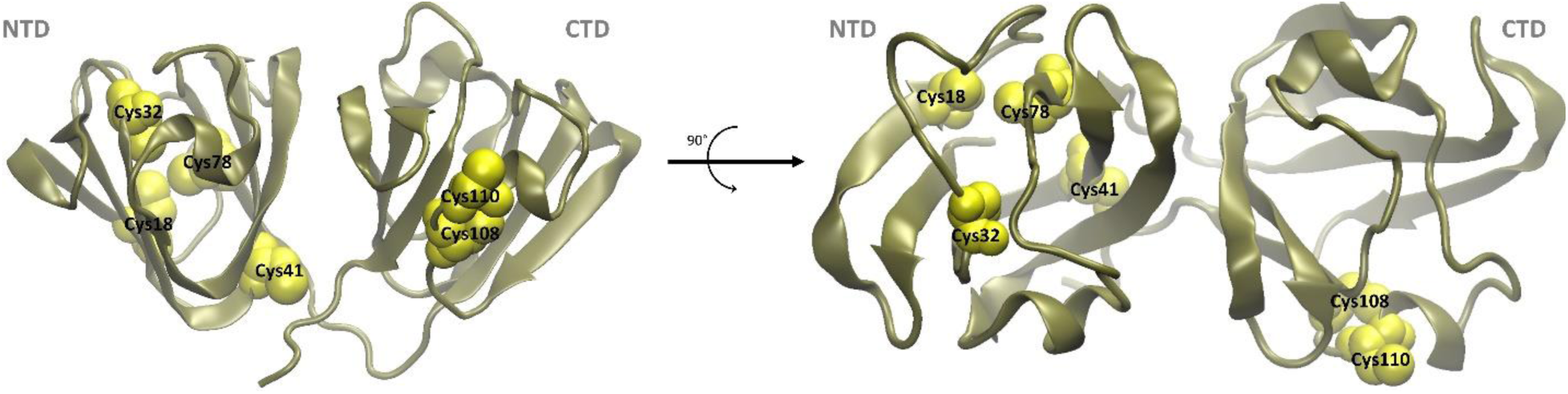
Locations of the Cys residues in the native structure of HγD (48)) (PDB ID 1HK0. By convention, we are using the PDB residue numbering here and throughout this paper.

As we detail below, the Cys>Ser substitutions caused a remarkably large decrease in chemical stability. Furthermore, dimers of natively folded molecules of this variant, held together by a disulfide at the C-terminal redox-active site (the only Cys residues we did not replace), revealed an unexpected aggregation pathway. This pathway led to similarly amorphous aggregates, and on a similar time scale, as the non-native disulfide-driven aggregates we have reported previously (29, 30, 41, 43, 44). However, this aggregation pathway required pre-existing dimers of natively folded molecules, and its concentration dependence was close to linear. Thus, the key rate-limiting step was intramolecular: misfolding of the dimer of natively folded molecules to an altered, aggregation-prone dimer. The NCA/T dimer showed a less severe, but qualitatively similar, phenotype. In the context of prior findings, our results suggest that many distinct perturbations to the NTD of HγD – whether mutational or chemical – can lead to comparable outcomes. If so, the misfolding process that occurs in the disulfide-bridged dimer of natively folded molecules, and the resulting misfolded dimer itself, could be targets for future pharmacological prevention or intervention for age-onset cataract.

## Results

### Cys-depleted variants are strongly destabilized relative to the WT

We expressed tag-free NCS and NCA/T HγD protein recombinantly in *E. coli* and purified them using ion exchange, hydrophobic interaction chromatography, and size exclusion, as described in Experimental Procedures. **Figure 2** shows that this procedure resulted in highly pure protein samples with minimal evidence of contamination or degradation. The WT protein was purified similarly (see Experimental Procedures). Short-term storage of samples was at +4 °C, and long-term storage was at −80 °C. Dimers formed spontaneously during storage at +4 °C over several days; addition of a mild oxidant (1% dimethyl sulfoxide, DMSO) to the storage buffer appeared to accelerate this process. Dimers and monomers were cleanly separated on an analytical/semi-preparative size exclusion column (**Figure 2A**). Electrospray mass spectrometry (**Supplementary Table 1**) confirmed that the dimer had a molecular mass of 2x(Monomer) – 2 Da., consistent with a single disulfide bridge per dimer. The dimers were essentially completely reverted to monomers under reducing SDS-PAGE (**Figure 2B**), also consistent with dimerization due to a disulfide. **Figure 1** shows the location of the only solvent-accessible Cys residue in these constructs, Cys110. All four N-terminal Cys residues were replaced in these constructs. We therefore conclude that the dimers were held together by a single disulfide at their C-terminal domains. This bond is probably C110-C110, as has been observed for WT HγD previously (34). This contrasts with the C24-C24 disulfide-bridged dimer observed in human γS-crystallin (HγS) (31, 49). We have previously noted that γ-crystallins tend to have solvent-exposed Cys residues in one domain or the other, but not both; otherwise garlands of crosslinked natively folded proteins would scatter light in the healthy aged lens (35). Distinct dimerization sites could have distinct consequences in HγD vs. HγS.

**Figure 2.**
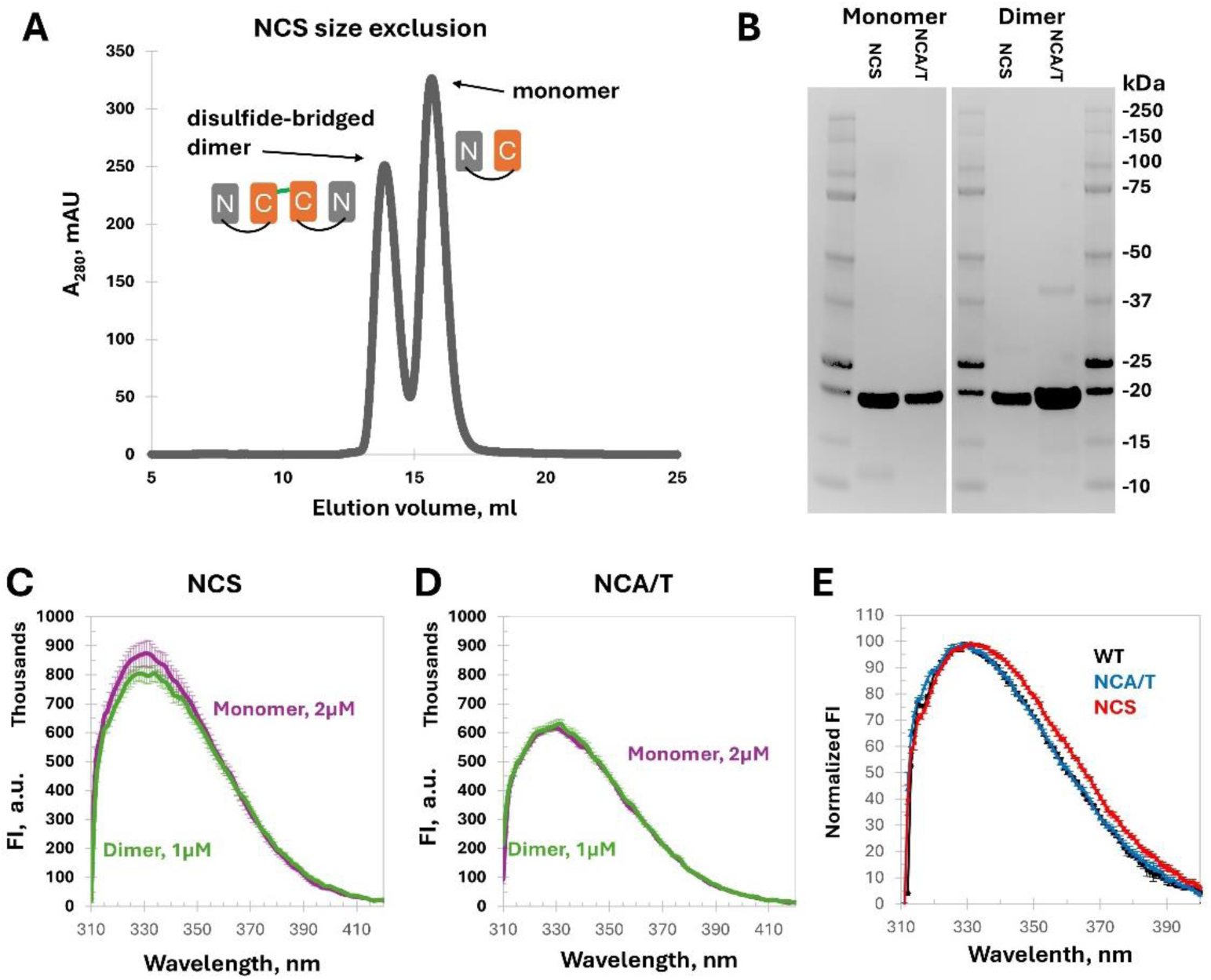
Biochemical characterization of monomeric and dimeric HγD-crystallin variants. (A) Size-exclusion chromatogram showing well-resolved monomer and covalent dimer peaks for the NCS variant. (B) Reducing SDS-PAGE of purified samples confirming high purity and the expected apparent molecular masses (∼20 kDa) for both the monomeric and the dimeric samples; the minor dimer band in NCA/T dimer is likely due to incomplete reduction. (C) Intrinsic tryptophan fluorescence spectra of NCS monomer (2 μM, *purple*) and NCS dimer (1 μM, *green*), without denaturant, excited at 280 nm and recorded from 310–420 nm. (D) Intrinsic tryptophan fluorescence spectra of the NCA/T variant monomer (2 μM, *purple*) and dimer (1 μM, *green*) collected under the same conditions. (E) Normalized fluorescence emission spectra were identical for WT and NCA/T but slightly red-shifted for NCS. Error bars for (C) and (D) are standard errors of three measurement replicates; those for (E) are standard errors of three replicate protein batches.

We observed very little difference between the intrinsic Trp fluorescence spectra of monomers and dimers of the NCS variant (**Figure 2C**). We did not observe any difference in intrinsic Trp fluorescence between the monomers and dimers of the NCA/T variant (**Figure 2D**). In **Figure 2C,D**, all monomers were measured at 2 µM and dimers at 1 µM, as measured by UV absorbance. Thus, the total concentration of Trp fluorophores was matched between monomer and dimer samples, as well as between the NCS and NCA/T samples, which were all measured side by side in the same 96-well plate. This experimental design made any differences in fluorescence intensity between monomers and dimers of a given protein as rigorously interpretable as possible by minimizing noise due to batch/batch variation, the instrument, or the assay vessel. The buried Trp pairs are sensitive reporters of HγD conformation, so the lack of either a spectral shift or even any significant difference in fluorescence emission intensity indicates the dimers were composed of natively folded molecules.

While this experiment rigorously compared monomers to dimers for each variant, the apparent difference in fluorescence intensity *between* variants is not rigorously interpretable due to noise in measuring protein stock concentrations and possible batch-to-batch variation in protein and buffer preps. However, internally normalized fluorescence spectra revealed a small but clear red shift in the emission of the NCS variant compared to NCA/T and the WT (**Figure 2E**). This is the origin of the slightly higher FI360/320 ratio at 0 denaturant in NCS compared to WT and NCA/T, as shown in **Figure 3**. This red shift indicates a slightly more hydrophilic environment, at least on average, of one or both Trp residues within the NTD of the NCS construct compared to the others.

**Figure 3:**
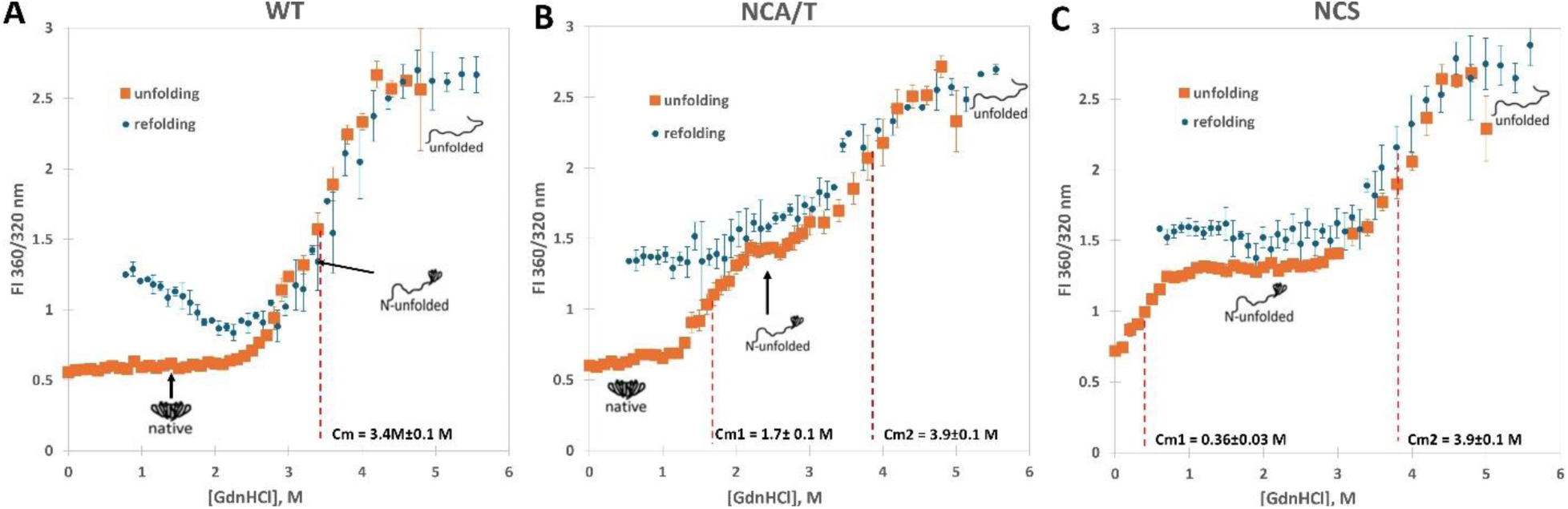
Thermodynamic stability by chemical unfolding and refolding. HγD WT Monomer (A), NCS Monomer (B), and NCA/T Monomer (C) were equilibrated with a series of guanidinium chloride (GdnHCl) concentrations and monitored by the fluorescence intensity ratio FI360nm/320nm (53). Orange squares indicate unfolding experiments (native protein directly incubated at the indicated [GdnHCl]); blue circles indicate refolding experiments (protein first fully denatured at high [GdnHCl], then diluted to the indicated [GdnHCl] and equilibrated). Error bars represent the standard errors of the mean (S.E.M.) of 3-6 replicates. Cm values, reported only for unfolding curves, represent the means and S.E.M. obtained by fitting each replicate individually to two-state (WT) or three-state (variants) Boltzmann sigmoids using Origin Pro®.

Equilibrium chemical denaturation experiments (**Figure 3**) showed pronounced three-state folding for both Cys-depleted variants. This is attributable to destabilization of the NTD, since all mutations were in that domain. Sigmoid fits yielded transition midpoint [GdnHCl] values (C_m_) of 0.36 ±0.03 M and 3.9 ±0.1 M for the NCS variant and 1.7 ±0.1 M and 3.9 ±0.1 M for NCA/T. The WT protein exhibited a very slight plateau in the equilibrium unfolding curve, consistent with the literature (50-52). However, only a single overall C_m_ of 3.4 ±0.1 M could be fitted. These results do not imply that the variants’ CTDs are more stable than the WT, simply that the WT C_m_ is a convolution of its NTD and CTD unfolding transitions. The degree of destabilization of the NCS variant was comparable to what we have previously observed for the W42Q variant (41) – a surprising effect from such a conservative set of amino acid substitutions: Cys>Ser is only a one-atom replacement, and the chemical properties of the side chains are considered similar. The NCA/T variant was less destabilized, despite the substitutions being somewhat less conservative; still, its loss of chemical stability was comparable to cataract-associated V75D variant (53).

For both NCS and NCA/T, the transition from FI360/320 ≈0.5 to FI360/320 ≈1.5 corresponds to the unfolding of the N-terminal domain, while the transition from FI360/320 ≈1.5 to FI360/320 ≈2.5 corresponds to the unfolding of the C-terminal domain. In the WT protein, the two transitions are so close to each other as to be essentially indistinguishable, consistent with prior literature reports (41, 50, 54-56). Kosinski-Collins *et al*. (54) reported Cm of ∼2.7 M for the WT protein; Vendra *et al.* reported 2.81 M (55); and most recently Volz *et al.* reported 2.98 ± 0.12 M (56). Our Cm value of 3.4 ± 0.1 M is therefore slightly above the range of the prior reports. This could be due to methodological differences; for example, all the above-mentioned studies measured Trp fluorescence using cuvette-based fluorometers with λ_ex._ = 295 nm; ours may be the first study to measure all fluorescence spectra in 96-well plates in a fluorescence plate reader, which required us to use λ_ex._ = 280 nm instead. We also cannot rule out minor differences due to reagents used, e.g., guanidinium chloride purchased as powder vs. as a concentrated solution. However, all the procedures and reagents in our study were the same for the WT and variant protein measurements.

For the WT protein, unfolding and refolding were fully reversible, with a minor cusp upward in the FI360/320 ratio attributable to aggregation competing with refolding upon dilution from high to very low denaturant concentrations, as previously observed (53, 54). The NCS variant showed a slight hysteresis at the plateau, but we could not reach sufficiently low denaturant concentrations to observe any refolding of its N-terminal domain. The NCA/T variant showed the strongest hysteresis in our experiment: its NTD practically did not refold even at guanidinium concentrations well below the unfolding transition. This hysteresis likely indicates kinetic trapping of the NTD in a misfolded monomeric or oligomeric state during refolding. Early studies using atomic force microscopy have found off-pathway oligomers formed during refolding of the WT protein from denaturant, including apparent small annular structures (54). Future work might similarly uncover the nature of the non-refolding-competent species in these HγD variants.

### Dimerization of natively folded molecules drives amorphous aggregation

Serendipitously, we found that the dimeric samples of both variants aggregated under mild conditions (**Figure 4B,D**), whereas the corresponding monomeric samples did not (**Figure 4A,C**). NCS dimers aggregated even under physiological conditions (the “0.0” trace in **Figure 4D**): 37 °C and pH 6.7, similar to the reported cytosolic pH in the lens core (57). Addition of low concentrations of denaturant (guanidinium) strongly promoted aggregation of the NCS and induced aggregation of NCA/T, as well.

**Figure 4.**
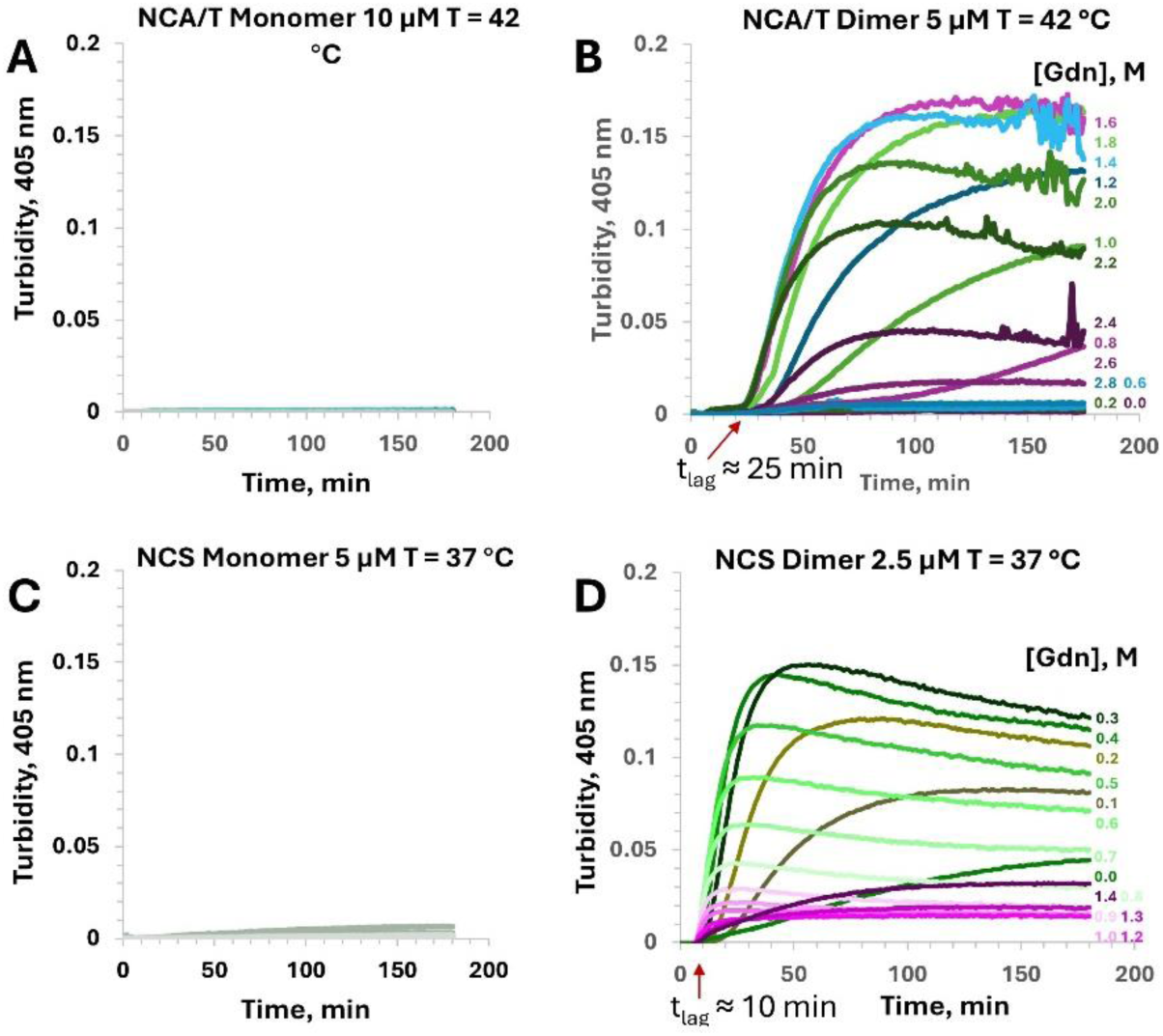
Aggregation of HγD NCA/T and NCS dimers. (A) NCA/T monomer (10 µM, 42 °C), (B) NCA/T dimer (5 µM, 42 °C), (C) NCS monomer (5 µM, 37 °C), and (D) NCS dimer (2.5 µM, 37 °C) were incubated in PIPES pH 6.7 buffer in the presence of increasing concentrations of GdnHCl, as indicated. Turbidity at 405 nm was monitored over 3 h, and representative time courses are shown for each condition. The concentration of monomeric samples was double that of the dimeric samples to ensure the same total number of protein subunits (i.e., the same mass concentration). Figure 5 emphasizes the overall patterns of how aggregation rate changes with [Gdn].

Disulfide crosslinking alone cannot explain this aggregation process: the aggregation-prone NTDs lack Cys residues and cannot form disulfides, while the CTDs can form disulfides but are wild-type and not prone to aggregation. Therefore, we hypothesized that aggregation requires non-native, non-covalent interactions formed by the misfolded NTDs, as explained below.

At higher [GdnHCl], the turbidity traces in **Figure 4B,D** show a rapid rise followed by a slow decrease. Two likely hypotheses could explain this decrease. The first is aggregate condensation, as previously reported (44): as aggregates become denser and more globular, their light scattering cross-section decreases. The second is a kinetic competition between aggregation and unfolding, particularly at these higher concentrations of denaturant.

Strikingly, while low [GdnHCl] strongly promoted aggregation of the dimers, higher [GdnHCl] strongly suppressed it. Plots of the maximum aggregation rate as a function of denaturant accordingly were bell-shaped (**Figure 5**). The peak of the aggregation rate for each variant closely matched the C_m_ value of its first unfolding transition, i.e., the [GdnHCl] at which the folding and unfolding rates of the NTD were evenly balanced at equilibrium. As shown in **Figure 6**, unfolding and aggregation occur on comparable time scales in these variants. If aggregation requires partially unfolded N-terminal domains, then it may be reversed as the domains unfold more fully. We have also previously observed denaturant-induced partial dissociation of aggregates formed by the W42E and W42Q variants of HγD (41, 43).

**Figure 5.**
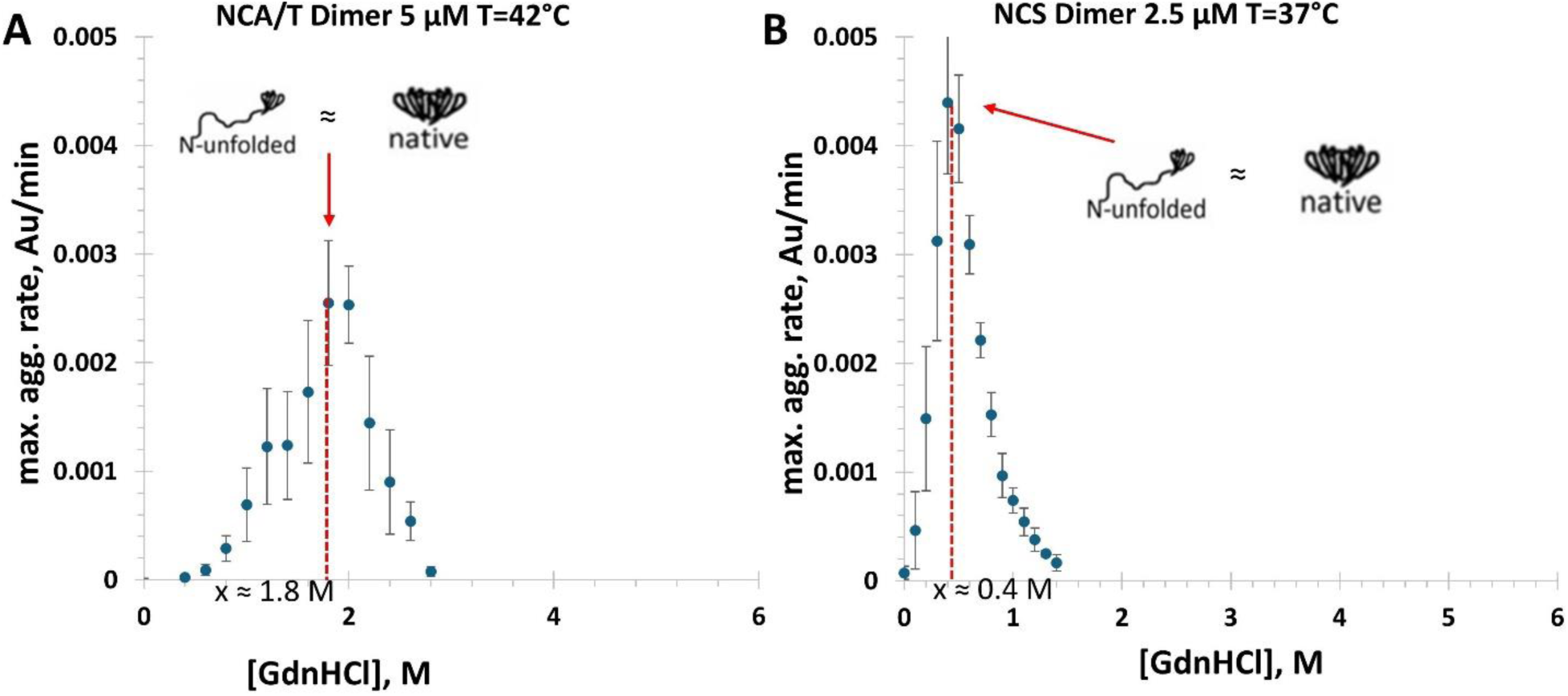
Guanidinium-dependent aggregation rates of γD-crystallin NCA/T and NCS dimers. (A) NCA/T dimer at 42 °C; (B) NCS dimer at 37 °C. For each condition, the maximal slope of the OD₄₀₅ vs. time trace was plotted as a function of GdnHCl concentration. These slopes were calculated from the curves shown in Figure 4, plus additional replicates. Error bars represent the standard deviation of measurements from three batches of each protein. Notably, the peaks of the bell-shaped curves were a close match to the C_m_ values of the respective NTD unfolding transitions, i.e., the aggregation rate was maximal when the concentration of folded and unfolded NTD’s was approximately equal.

**Figure 6.**
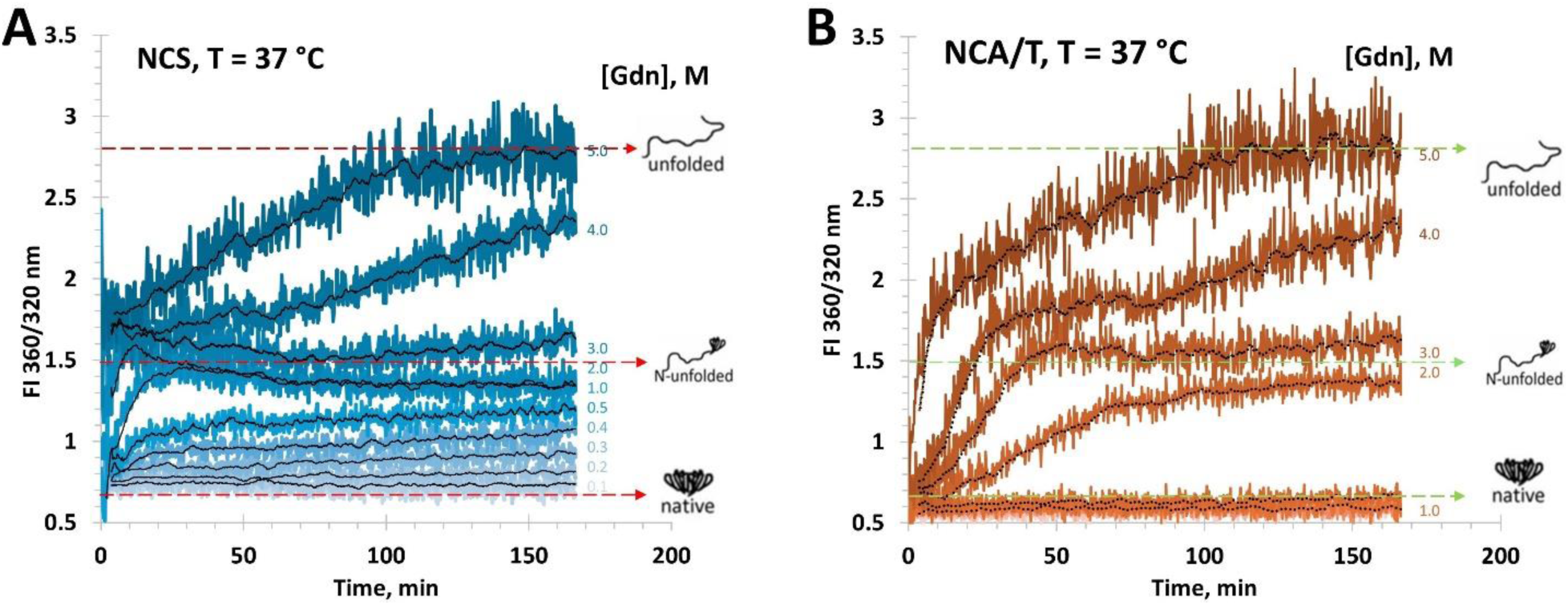
Time-resolved unfolding kinetics of NCS and NCA/T HγD-crystallin variants. NCS monomer (A) and NCA/T monomer (B) unfolding kinetics were monitored by the FI360nm/320nm ratio over 200 min at the indicated GdnHCl concentrations. For each denaturant concentration, the kinetic traces are shown in gradient color, and the central black line represents the mean trajectory (moving average) for that condition. Horizontal dashed lines mark the approximate FI360/320 values corresponding to the native baseline (“native”), the intermediate (“N-unfolded”) with the NTD unstructured but CTD still structured, and the fully unfolded state (“unfolded”), as inferred from equilibrium measurements in Figure 3.

HγD WT is known to have very slow unfolding and refolding rates (51), so we wondered whether the NCS and NCA/T dimers in the Gdn-promoted aggregation experiments in the present study were near equilibrium or only beginning to unfold. We used FI360nm/320nm measurements in the kinetic mode, correcting for the ∼30 s. dead time of plate loading (**Figure 6**). To avoid confounding effects of aggregation, we used the monomeric proteins for these measurements, as for the equilibrium measurements in **Figure 3**. We observed that, in the [Gdn] range of **Figure 5B**, NCS equilibrated within minutes and was therefore already near equilibrium at the peak aggregation rate reported in that figure. In the case of NCA/T, however, the sample was no more than 50% of the way to equilibrium by the t_lag_ (∼25 min) of the aggregation assay (**Figure 4C**). This likely explains why the two highest aggregation rates in **Figure 5A** occurred at 1.8 M and 2.0 M GdnHCl – slightly above the 1.7 ±0.1 M C_m_ for the NTD melting transition in **Figure 3C**.

HγD and its variants have been shown to form various types of aggregates under various conditions. These include reversible native-state aggregates (58), amyloid aggregates (54, 59), mixed amyloid and amorphous aggregates (60), metal-bridged aggregates (61), UV-induced amorphous aggregates (62), and amorphous aggregates trapped by intramolecular disulfides (41, 43, 44). The most physiologically relevant conditions appear to give rise to amorphous aggregates. Transmission electron microscopy (**Figure 7**) showed that the NCS dimer aggregates in the present study were amorphous and qualitatively similar to the UV-induced WT aggregates (62) and the aggregates of the W42Q variant that mimics UV-catalyzed Trp oxidation (44).

**Figure 7.**
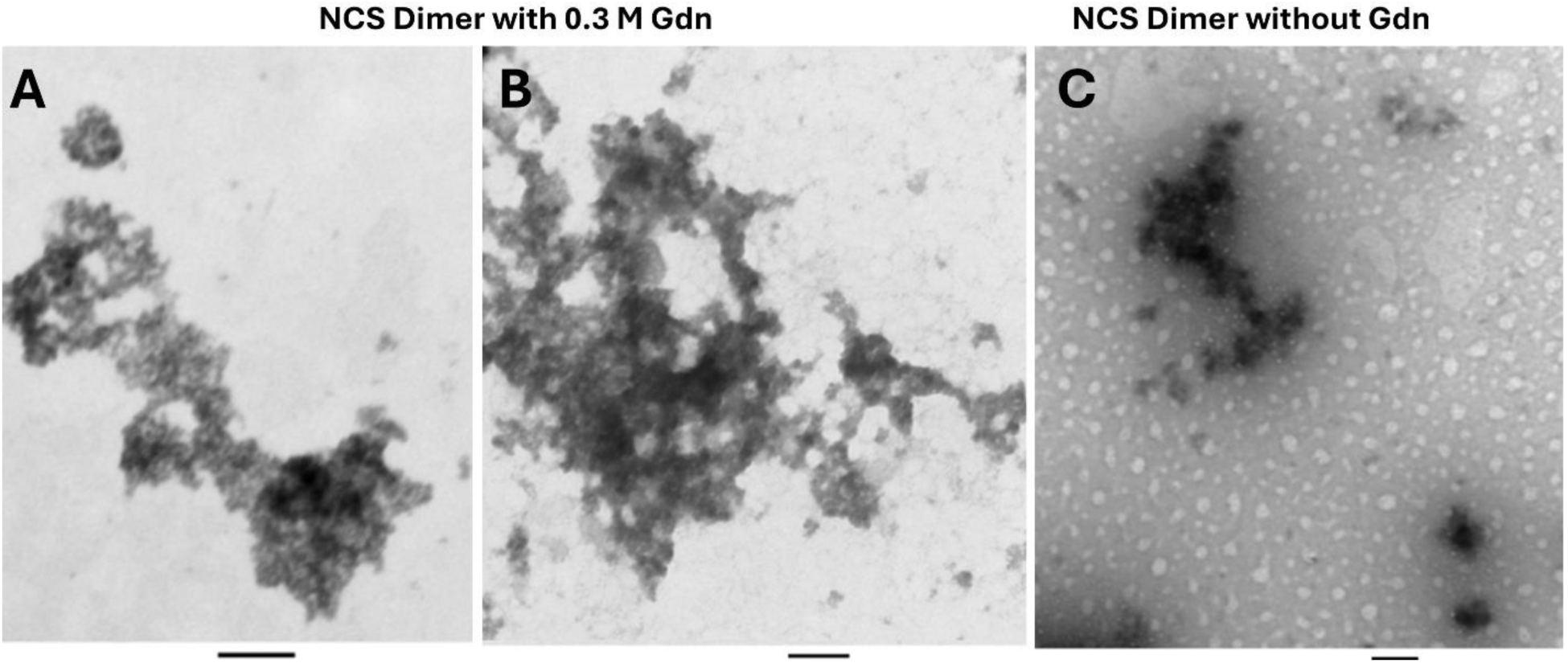
Transmission electron microscopy (TEM) images of HγD NCS dimer aggregates. Representative TEM micrographs of aggregates formed by the NCS dimer under the aggregating conditions used in this study, with and without 0.3 M GdnHCl, as indicated. The scale bar at the bottom of each image corresponds to 100 nm.

### Intramolecular misfolding within the dimer of natively folded molecules determines the aggregation rate

The observation that maximum aggregation rate occurs when the NTD is near the midpoint of its equilibrium unfolding transition (**Figure 5**) is consistent with a model in which a misfolding event within the dimer is the rate-determining step for aggregation. That is, when both NTD’s within the dimer fluctuate rapidly between folded and unfolded states, there is a competition between refolding *in cis* back to the native state and refolding or misfolding *in trans* to form a domain-swapped or pseudo-domain swapped state. The latter is a state that reconstitutes *in trans*, not the native conformation, but an alternative or fold-switched state, as we have previously proposed for the W42Q variant (29).

Since the misfolding event within the dimer is intramolecular, the model makes a clear prediction: the aggregation rate should vary linearly with dimer concentration. This prediction is in sharp contrast with all our prior findings on the aggregation of HγD variants: in all those cases, the concentration dependence was approximately quadratic (41, 44, 63). We therefore measured the maximal aggregation rate over a 23-fold range of NCS dimer concentrations at 42 °C. Indeed, the concentration dependence was approximately linear (**Figure 8C**). A straight-line fit to the data without inositol was nearly as good as the power-law fit shown in the figure (R^2^ = 0.94 vs. 0.96), while in the presence of *myo-*inositol the straight-line fit was better (R^2^ = 0.98 vs. 0.92). We also measured the concentration dependence of NCS dimer aggregation over a 23-fold range in the presence of 0.3 M GdnHCl at 37 °C and compared it with aggregation at 42 °C without GdnHCl: linear concentration dependences were observed in both cases (**Figure 8E**). Meanwhile, the lag time of aggregation showed little or no concentration dependence over this range (**Figure 8D**), which also contrasts with our prior observations with W42Q and other variants (63). We conclude that the rate-determining step for aggregation in all cases is formation of misfolded dimers; in the prior studies, this required two misfolded monomers to come together (a bimolecular interaction), while in the present study, it requires only the transition of a natively folded dimer to a misfolded dimer (a unimolecular event).

**Figure 8:**
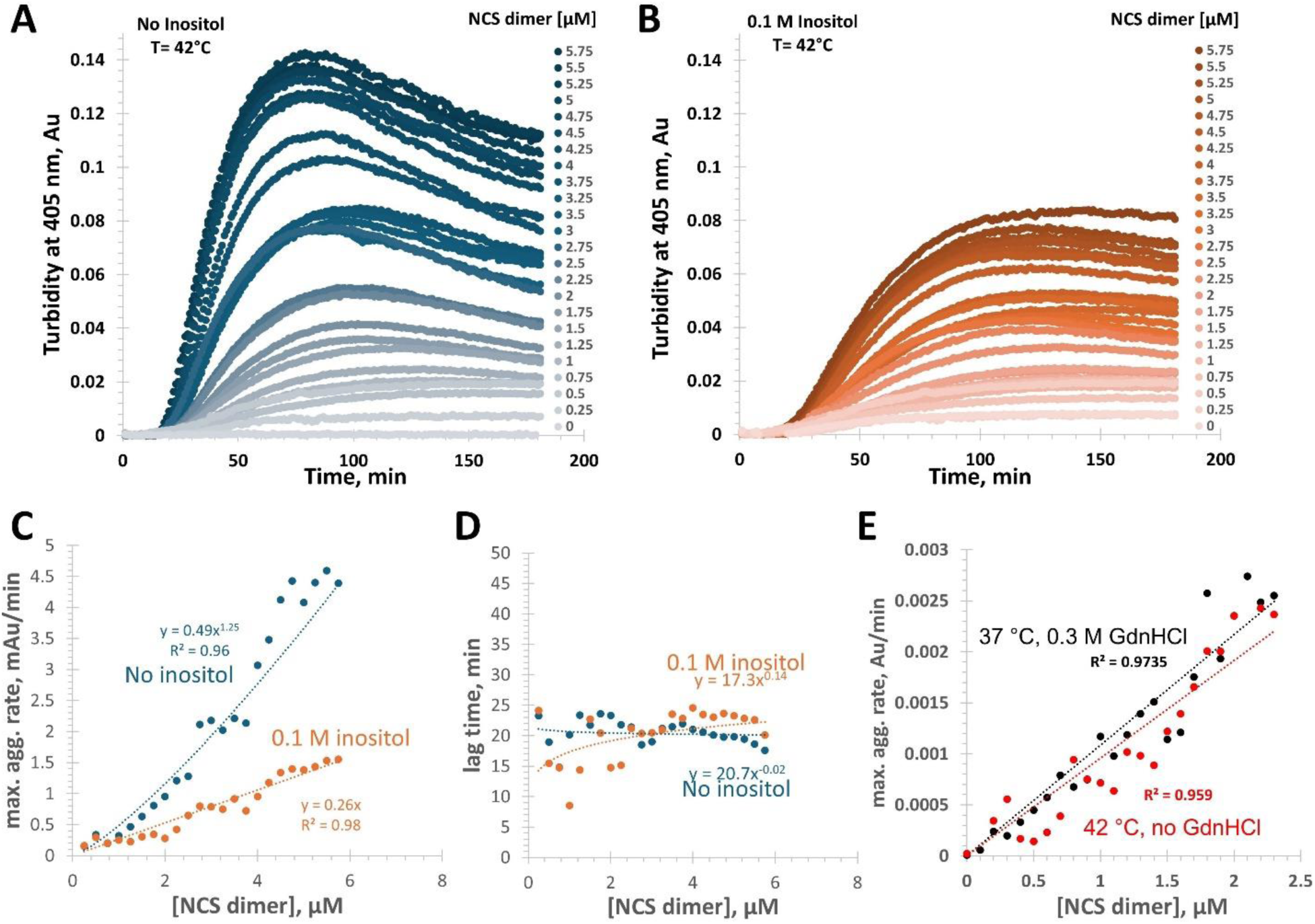
Concentration dependences of NCS dimer aggregation rates. The NCS dimer sample was aggregated at 42 °C without any chemical denaturant at the indicated concentrations with (A) and without (B) a physiologically relevant level of 0.1 M *myo*-inositol (44). (C) The maximal aggregation rate depended near-linearly (∼x^1.25^) on [NCS dimer] without *myo*-inositol and fully linearly with *myo-*inositol. Given the noise in the data, the concentration dependence should be considered approximately linear in both cases. (D) The apparent lag times of aggregation showed little or no dependence on protein concentration. (E) In an independent experiment, the concentration dependence of the maximal aggregation rate was approximately linear both with and without 0.3 M GdnHCl; linear fits are presented as eye guides here.

### The native chemical chaperone *myo-*inositol inhibits the rate-determining misfolding step

The healthy human eye lens contains an estimated ∼80 mM of *myo*-inositol in the free water fraction (44), and the lenses of other primates may be even more inositol-rich (64). We recently reported that *myo*-inositol at physiologically relevant concentrations can act as a chemical chaperone, suppressing aggregation of HγD variants destabilized by mutation or truncation (44). *Myo*-inositol also clearly suppressed aggregation of the NCS dimer in the present study (**Figure 8B,C**). To further quantify this effect, we measured aggregation of the NCS dimer with a 23-fold range of [*myo*-inositol]. The aggregation suppression was strong. The data appeared to be better fit by an exponential decay function than a linear function, but we do not draw mechanistic conclusions from this. (**Figure 9**). Remarkably, even the lowest concentrations of *myo*-inositol assayed here yielded significant aggregation suppression – unlike our prior observations with the oxidative aggregation of W42Q, W130Q, or NTD-only variants (44).

**Figure 9:**
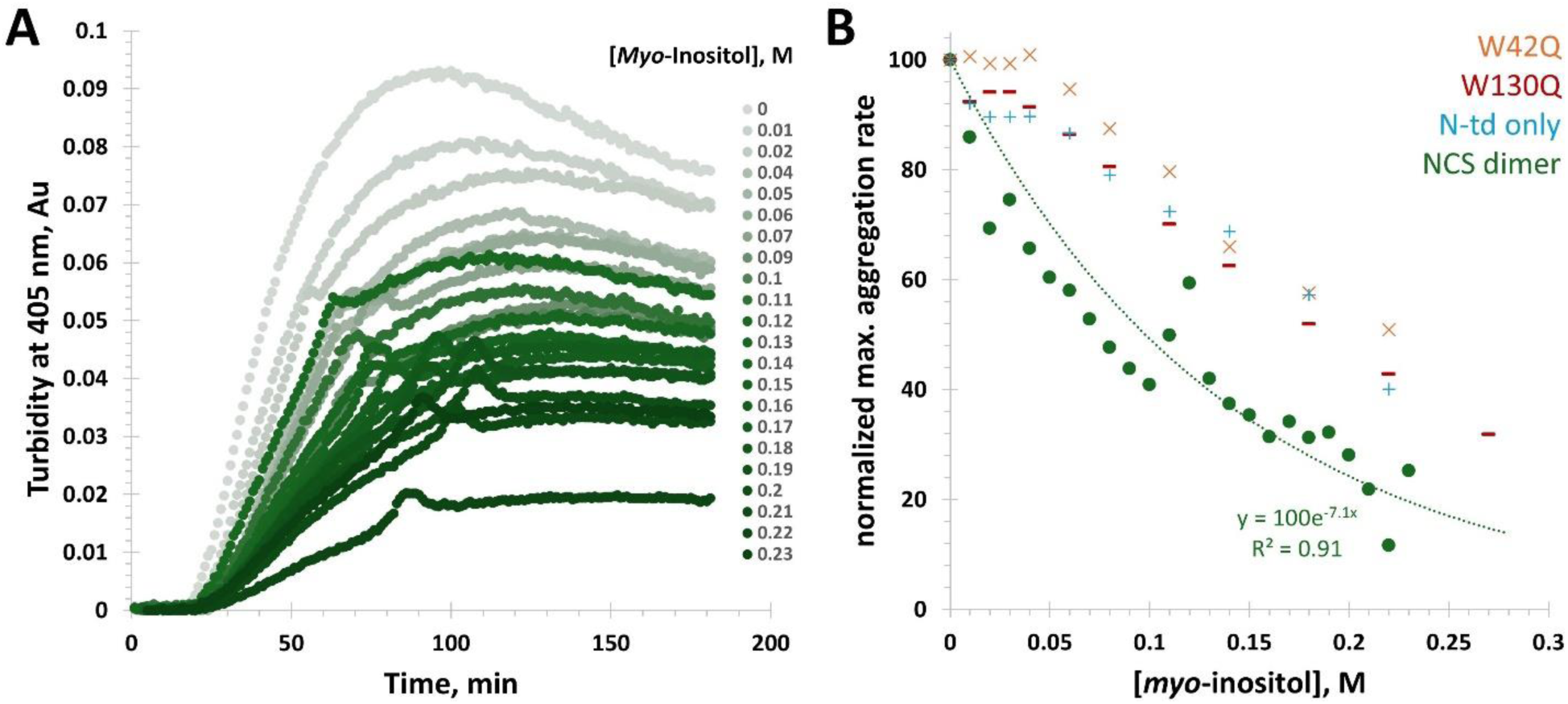
Suppression of NCS dimer aggregation by the lens metabolite *myo*-inositol. (A) Solution turbidity traces for 4 μM NCS dimer at 42 °C with varying [*myo-*inositol] as indicated. (B) Suppression of the maximal aggregation rate (expressed as % of the no-inositol condition) as a function of [*myo-*inositol] was notably stronger for the NCS dimer than for previously reported oxidatively aggregating HγD variants and fit an exponential decay function. Data for the W42Q, W130Q, and NTD-only samples are from ref. (44).

## Discussion

We have shown that removal of the native buried Cys residues from the core of the HγD NTD results in major loss of thermodynamic stability and an unexpected aggregation pathway initiated by intramolecular misfolding within a dimer of natively folded molecules. Prior research found that replacing all Cys residues in HγD with Ser resulted in significant loss of thermostability, with the T_m_ decreasing from ∼82 °C for WT to ∼60 °C for the all-Ser variant (65). By comparison, the first melting transition of the highly aggregation-prone W42Q variant is ∼55 °C (29). Thus, the chemical and thermal denaturation data agree: the NTD is destabilized by the four seemingly conservative Cys>Ser substitutions as by the seemingly much more drastic W42Q substitution. The former may be a case of “death by four cuts” for the HγD NTD.

The W42Q substitution mimics the effect of Trp oxidation to kynurenine or a similar hydrophilic product (41, 43). The Cys>Ser substitutions may mimic the mildest form of Cys oxidation: conversion to sulfenic acid (66). Such a modification is typically too unstable to detect proteomically, but UV exposure has been shown to result in oxidation of at least one HγD NTD residue to the highest (and stable) oxidation state: sulfonate (67). Conversion of Cys to sulfonate might be mimicked by Cys>Asp substitutions, but we have not attempted this due to the already strong destabilization caused by Cys>Ser.

We have previously reported that aggregation of cataract-associated HγD variants under physiological conditions requires non-native disulfides (29, 41). The findings of the present study indicate that aggregation can also proceed without any disulfides in the NTD, provided that a CTD disulfide first brings two natively folded HγD molecules together to form a dimer. We observed that the misfolding-limited aggregation rate was maximal under conditions that correspond to the midpoint of the NTD equilibrium unfolding curve. Yet, dimers prior to the aggregation experiment had Trp fluorescence spectra identical to the monomers. Thus, our working conceptual model is that dimerization ensures very high effective concentration of natively folded HγD molecules, so that, when the two NTD’s unfold and refold in parallel within the dimer, they may intercalate to form a misfolded dimer instead of returning to their native conformations. Future studies should investigate whether this is an example of domain swapping (68-71) or of a pseudo-domain swapping mechanism akin to what we have previously proposed for the W42Q variant (29).

In general, the accumulating evidence that many distinct perturbations to the HγD NTD all lead to misfolding and aggregation suggest a model of convergent misfolding pathways. That is, HγD has one class of highest-likelihood altered states, and any perturbation that raises the free energy of the native NTD conformation makes it more likely that the protein will visit the altered state that serves as the precursor to aggregate formation (**Figure 10**). This perturbation could be mutational or, equivalently, post-translational, such as deamidation, racemization, truncation, sulfonation, etc. In this proposed model, disulfide-bridged dimerization of natively folded molecules does not directly destabilize the native state, but it favors the altered state and facilitates the transition. Conversely, the effect of *myo-*inositol may be conceptualized as partially reverting this free energy landscape toward what it had been for the monomer. This hypothesis is consistent with the prior findings that *myo*-inositol has no discernible binding site on the native structure, nor any significant effect on the stability of the native state, yet it has significant effects on the unfolding pathway, making it more cooperative (44). In the exceptionally well-controlled system we now report, where aggregation occurs only after a misfolding event within the dimer, the origin of the increased cooperativity seems straightforward. We propose that *myo*-inositol limits interactions between the subunits of the dimer as the NTDs of those subunits fluctuate between folded and unfolded conformations. It has long been known in the protein folding field that rugged landscapes of protein unfolding frequently result from misfolded dimers or oligomers (72, 73). Studies of multidomain and repeat proteins have concluded that intramolecular misfolding plays a major role, leading to very rugged free energy landscapes (74, 75). Intramolecular misfolding, driven by mislocalization of the N-terminal β-hairpin, has been observed in engineered HγD repeats (polyproteins) (76) and in atomistic simulations (29). In the present study, disulfide-bridged dimerization may serve the same function: bringing the subunits close enough together to turn rare intermolecular misfolding into a frequent intramolecular (i.e., within the covalent dimer) event.

**Fig. 10:**
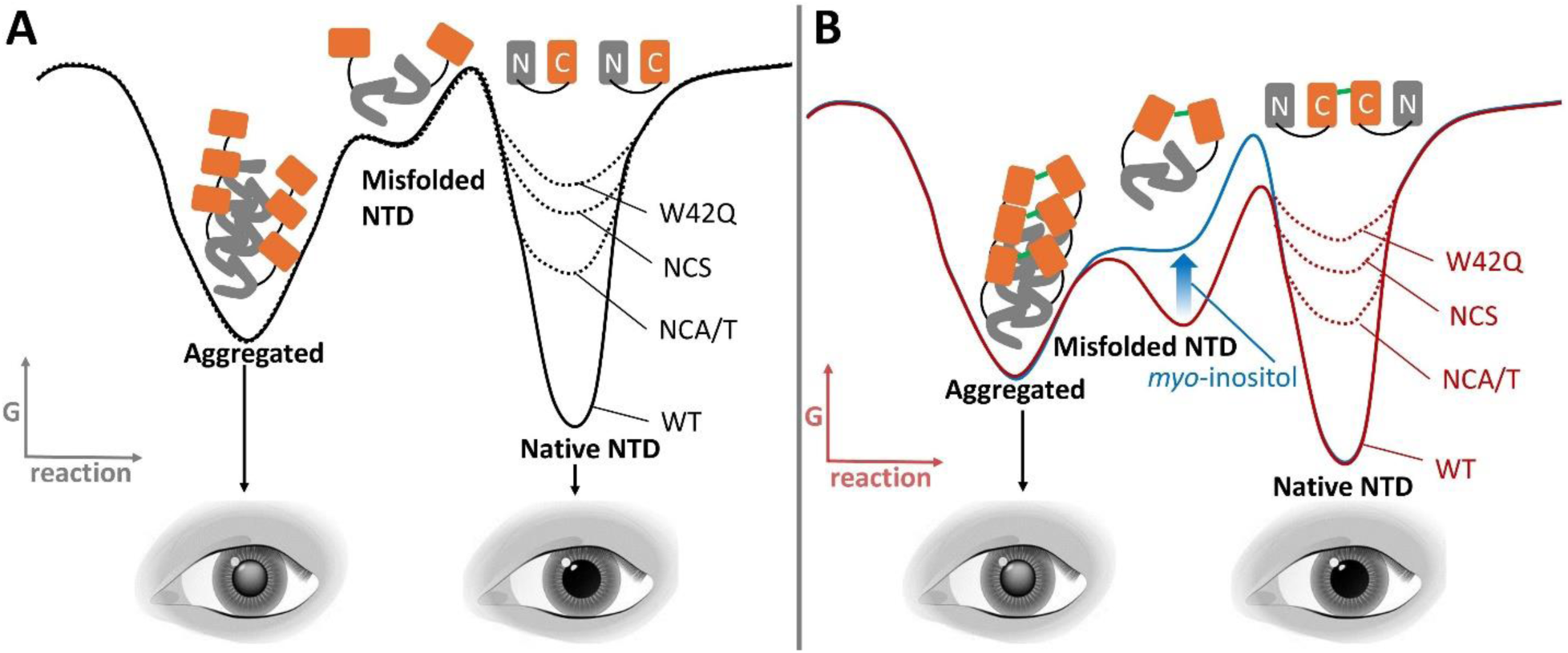
Working model of convergent misfolding pathways in HγD. This schematic depicts the landscape before (A) and after (B) disulfide-bridged dimerization. Dimerization of natively folded molecules facilitates formation of altered NTD conformers (*red curve*), while presence of *myo-*inositol partially restores it (*blue arrow* and *blue curve*). Various mutations or post-translational modifications destabilize the native conformer and thereby drive the protein toward the altered and aggregated conformations.

The concentration dependence of the maximal aggregation rate was quadratic in all our previous reports, but in the present system it is linear. This observation, too, is explained by the effect of dimerization: it coverts a bimolecular rate-limiting step (misfolding of two interacting monomers) into a unimolecular one (conformational rearrangement within a covalent dimer).

Dimerization often stabilizes protein molecules. In fact, while all human γ-crystallins are natively monomeric, all human β-crystallins are natively dimers or tetramers (77). HγD forms dimers of natively folded molecules readily, due to the solvent-exposed Cys110 residue in its stable C-terminal domain (34). This residue is subject to some biological regulation, being post-translationally blocked (methylated) in approximately half of HγD molecules in the lens (78). Its continued availability in the remaining HγD molecules may be due to the usefulness of the C108-C110 pair as an oxidoreductase site. Indeed, as the lens loses its native glutathione buffer with age, HγD and other highly abundant and long-lived lens γ-crystallins likely take over the redox buffering function (30). A previous study reported a moderate increase in oxidative aggregation propensity in a γS-crystallin dimer (79). However, we have serendipitously found an aggregation process that crucially depends on a misfolding event that occurs only in the context of the disulfide-bridged dimer of natively folded molecules.

*Myo*-inositol, a highly abundant lens metabolite, likely serves as a native aggregation inhibitor of lens crystallin aggregation (44), and loss of *myo*-inositol correlates with cataract disease (80). We found that this compound inhibits the aggregation of NCS dimers even more strongly than that of the constructs we have previously reported (44). There could be two reasons for this. First, the misfolded conformers are no longer locked by long-lived non-native disulfide bonds. The weak non-covalent interactions of *myo*-inositol with HγD cannot compete with disulfide-trapped misfolding, but they may better compete with the non-covalent interactions that drive misfolding in the present study. Second, it is possible that, under the mildly denaturing aggregation-permissive conditions, the C110-C110 disulfides that bridge dimers may slowly convert to intramolecular C108-C110 disulfides. This would break the dimer and thus also revert the free energy landscape in **Figure 10** back to its “monomeric” shape. However, the dimers are very stable in storage, and we have not observed any spontaneous monomerization.

Our study sheds some light on the mechanism of aggregation suppression. *Myo*-inositol suppresses NCS dimer aggregation even at the lowest assayed concentrations, resulting in an apparent exponential-decay dependence of aggregation rate on [*myo*-inositol]. Prior findings with oxidative aggregation yielded a more sigmoid fit, with only modest effects observed below ∼50 mM *myo*-inositol (**Figure 9**). In other words, aggregation driven by non-native disulfides within the NTD escapes suppression by low [*myo*-inositol] more efficiently than that reported here. We hypothesize that *myo*-inositol inhibits the earliest stages of misfolding, which, for the variants reported here, occur within the dimer and are reversible. By contrast, W42Q under fully oxidizing conditions is likely trapped by a non-native disulfide within the NTD prior to the rate-determining dimerization step (44). Dimers of natively folded molecules of WT HγD are likely common in the aging lens (4, 34), and we propose that many forms of oxidative damage can destabilize the NTD’s within such dimers. If so, many forms of damage may facilitate conversion of dimers of natively folded HγD molecules into dimers of altered molecules. *Myo*-inositol may contribute to maintaining lens transparency at least in part by suppressing misfolding within dimers of natively folded HγD.

Our study was sparked by an evolutionary question: Why have γ-crystallins not evolved to be Cys-depleted, like the α-crystallins (35)? One possible answer based on our results is that HγD is “addicted” to its buried Cys residues. Single-nucleotide changes, the most frequent mutation type, can convert a Cys codon into a Trp, Tyr, Phe, Arg, Ser, Gly, or Stop codon. Ser is considered the most conservative among these substitutions, yet we have shown that replacing the four NTD Cys codons with Ser codons resulted in a severely destabilized protein. Ala/Thr replacements were less severe, but nonetheless substantially decreased stability. However, the increasing success of computational algorithms in designing highly thermodynamically stable Cys-free proteins, along with the overall modest conservation of Cys residues among the βγ-crystallins (35), suggests that highly stable Cys-depleted γ-crystallins can probably be designed or evolved. A more nuanced possibility is antagonistic pleiotropy. The high Cys content of γ-crystallins may maximize stability and refractive function of the lens proteome early in life, at the cost of disulfide-driven misfolding and aggregation later in life (35). These costs may be partially compensated by other adaptations, such as the crystallin redox buffer (30), the native chemical chaperone *myo*-inositol, and perhaps even by UV-catalyzed scission of disulfide bonds (4, 66).

The unexpected yet remarkably controllable dimerization-driven misfolding and aggregation of the two Cys-depleted variants converts what had been an intermolecular misfolding process into a much more tractable intramolecular misfolding event. This misfolding process, and the structure of the resulting misfolded state, are now open to detailed investigation and may yield new potential targets for structure-based pharmacological prophylaxis or treatment of age-onset cataract.

## Experimental Procedures

### Plasmids and competent cells

Plasmids encoding human γD-crystallin (wild-type and the Cys-depleted variants) were stored at −20 °C, and chemically competent E. coli BL21(DE3) cells were stored at −80 °C. Before transformation, plasmids and competent cells were thawed on ice for 5–10 min.

### Transformation

For each transformation, 50–100 ng of plasmid DNA were added to 26 µL of competent BL21(DE3) cells in a pre-chilled 1.5 mL microcentrifuge tube. The mixture was gently flicked to mix and incubated on ice for 30 min. Cells were then heat-shocked at 43 °C for 45 s, immediately returned to ice for 5 min, and recovered by adding 500 µL of LB broth (without antibiotics). Cultures were incubated at 37 °C with shaking at 230 rpm for 45 min, then spread onto LB agar plates containing 100 µg/mL ampicillin and 30 µg/mL chloramphenicol. Plates were incubated overnight at 37 °C to obtain single colonies.

### Small-scale pre-cultures and glycerol stocks

A single colony was picked with a sterile pipette tip and inoculated into 6 mL of Terrific Broth (TB) supplemented with 100 µg/mL ampicillin and 30 μg/ml chloramphenicol. The culture was grown overnight at 30 °C with shaking at 230 rpm. To prepare glycerol stocks, 300 µL of the overnight culture were mixed with 100 µL of 60 % (v/v) glycerol in a cryovial, thoroughly mixed, and stored at −70 °C. These glycerol stocks were used as a source for subsequent pre-cultures to increase the reproducibility of protein expression.

### Large-scale autoinduction expression

For large-scale expression, 600 mL of Terrific Broth(TB) medium were prepared in 2 L conical flasks by dissolving 28.56 g TB powder in 600 mL deionized water and sterilizing by autoclaving. After cooling to room temperature, glycerol, 0.05 % (w/v) glucose, 0.2 % (w/v) lactose, and 1 mM MgSO₄ were added aseptically to generate lactose autoinduction medium similar to that used previously (81, 82). Each flask was inoculated with 6 mL of overnight pre-culture. Cultures were incubated at 37 °C with shaking at 200 rpm until the OD₆₀₀ reached 0.6–0.8, at which point the temperature was reduced to 16 °C and incubation continued for 16–18 h to promote soluble protein expression. Aliquots (100 µL) were collected before and after the overnight induction for SDS-PAGE analysis of expression levels.

### Cell harvesting

Cells were harvested by centrifugation at 3750 rpm (∼4000 × g) for 1 h 20 min at 4 °C in 500 mL centrifuge bottles. The resulting cell pellets were resuspended in sample buffer and stored at −70 °C or −80 °C until purification.

### Purification of Human γD-Crystallin Variants

WT was purified as described (41). The purification of the variants was as follows.

### Cell lysis

Frozen cell pellets were thawed on ice and resuspended in ice-cold lysis buffer containing 10 mM PIPES, 50 mM NaCl, pH 6.7, lysozyme (100 μg/ml final), and one EDTA-containing mini protease inhibitor tablet per tube (cOmplete™-Mini, Pierce or Roche). Suspensions were gently mixed on a rocker at 4 °C for ∼15 min to allow complete dissolution of the inhibitor tablet and lysozyme action.

Cells were then disrupted by probe sonication (QSonica model Q500) on ice (40 % amplitude, 2 s on / 2 s off, total sonication time 5 min). The lysate was clarified by centrifugation at 40 000 × g for 30 min at 4 °C. The supernatant was carefully transferred to a fresh tube and centrifuged a second time at 40 000 × g for 30 min to ensure complete removal of insoluble debris. The resulting clarified supernatant was used as the starting material for hydrophobic interaction chromatography (HIC).

### Hydrophobic interaction chromatography (HIC)

To prepare the sample for HIC, solid ammonium sulfate or a concentrated ammonium sulfate stock solution was added over several minutes to the clarified lysate with gentle stirring until the final concentration reached 1.5 M. The sample was incubated on ice and then centrifuged at 40 000 × g for 20 min at 4 °C to remove any precipitated material. The resulting clear, high-salt supernatant was used for column loading.

HIC purifications were performed on an ÄKTA FPLC system with a column packed with approximately 10mls of Cytiva Phenyl Sepharose™ High Performance hydrophobic interaction column (HIC) medium. The column and system were equilibrated with binding (high-salt) buffer composed of 1.5 M ammonium sulfate, and 50 mM NaCl, and 10 mM PIPES, pH 6.7. A low-salt buffer (50 mM NaCl, 10 mM PIPES, pH 6.7) was prepared as the elution buffer.

After manual priming of the pumps and washing of the sample loop with the binding buffer, the HIC column was equilibrated with the binding buffer by running an equilibration method until approximately 60 mL of buffer had passed through the column. Up to 15 mL of High-salt protein sample was then loaded. Following sample application, the column was washed with 60 ml of binding buffer to remove unbound and weakly bound contaminants. HγD variants were eluted using a decreasing-salt gradient generated by mixing the binding and elution buffers, lowering the ammonium sulfate concentration from 1.5 M to 0 M over the course of 180 ml at a flow rate of 0.5 ml/min while maintaining 50 mM NaCl and 10 mM PIPES, pH 6.7. Elution was monitored by UV absorbance at 280 nm, and fractions were collected across the gradient, including the flow-through. Fractions containing major protein peaks were analyzed by SDS-PAGE, and those containing γD-crystallin of acceptable purity were pooled for desalting and further polishing.

### Desalting and buffer exchange

To remove ammonium sulfate and exchange the protein into size-exclusion chromatography (SEC) buffer, pooled HIC fractions were desalted on a HiPrep 26/10 Desalting column (53 mL bed volume, Sephadex G-25 Fine; Cytiva) connected to an ÄKTA GO chromatography system. The column was equilibrated with at least two column volumes of SEC buffer (10 mM PIPES, 150 mM NaCl, pH 6.7) at a flow rate of 5–10 mL/min, below the manufacturer’s recommended maximum flow rate (15 mL/min).

For each run, up to 15 mL of pooled HIC sample (≤1/3 of the column bed volume) were applied, in accordance with the recommended sample volume range for HiPrep 26/10 Desalting columns. Samples were loaded using SEC buffer, and elution was carried out isocratically in the same buffer. UV absorbance at 280 nm was monitored throughout the run. Early eluting fractions corresponding to the void volume peak (containing desalted protein) were collected, whereas later fractions containing small molecules and salts (including ammonium sulfate) were discarded.

### Size-exclusion chromatography (SEC)

Final polishing and isolation of monomeric γD variants were achieved by SEC on an ÄKTA Go system equipped with HiLoad^TM^ 26/600 Superdex^TM^ 75 pg column. Before sample injection, the column was equilibrated with SEC buffer (10 mM PIPES, 150 mM NaCl, pH 6.7), matching the buffer composition used during desalting. Desalted protein samples (typically up to 10 mL) were centrifuged at 3750 rpm for 20 minutes to remove any potential precipitates. For subsequent separation of covalent dimers from monomeric γD-crystallin, an additional SEC step was performed on a smaller Superdex^TM^ 75 Increase 10/300 GL column using the same buffer conditions.

Elution was monitored by absorbance at 280 nm, and fractions were collected automatically according to the absorbance profile. Fractions corresponding to the major monomeric peak were analyzed by SDS-PAGE. Fractions displaying a single band at the expected molecular weight of γD-crystallin were pooled as the final purified product and stored at 4°C until use, long-term storage requires keeping at −70°C.

### Thermodynamic Stability Measurements (Equilibrium Unfolding)

Thermodynamic stability of γD-crystallin variants was assessed by Gdn-induced equilibrium unfolding in a 96-well plate format (half area UV-Transparent Microplates). A series of wells with final Gdn concentrations spanning 0–5 M was prepared to probe the full unfolding transition.

Each well contained a total volume of 50 µL. Appropriate volumes of Gdn stock solutions were added to achieve the desired denaturant concentrations (e.g., 0, 0.1, 0.2, 0.3, 0.4 M up to 5 M). Protein was added from a concentrated stock such that the final protein concentration in each well was 1 µM. Buffer components were adjusted so that all wells contained identical concentrations of 0.1 mM EDTA, 1 mM DTT (to maintain a reducing environment and prevent unintended disulfide formation), 10 mM phosphate, and NaCl at a constant ionic strength. Water was added as needed to bring each well to the final volume of 50 µL.

After preparation, the plate was sealed and incubated at 37 °C for approximately 24 h to allow the protein to reach equilibrium at each Gdn concentration. Intrinsic tryptophan fluorescence was recorded on a SpectraMax® iD3 multimode microplate reader using an excitation wavelength of 280 nm and scanning emission from 310 to 420 nm. For each well, the fluorescence intensities at 320 nm and 360 nm were extracted, blank (no-protein) fluorescence was subtracted from all wells before computing FI₃₆₀/FI₃₂₀. The ratio FI₃₆₀/FI₃₂₀ was then calculated. Because unfolding red-shifts intrinsic Trp emission, this ratio was used as an operational measure of the fraction of unfolded protein. Plots of FI₃₆₀/FI₃₂₀ versus [Gdn] were used to generate equilibrium unfolding curves and to compare domain stability and unfolding cooperativity among the γD-crystallin variants. C_m_ values were determined by fitting each replicate curve individually to Botlzmann sigmoid two- or three-state curves using Origin Pro® and reported as mean ± S.E.M. for WT (n = 4), NCS (n = 6), and NCA/T (n = 6).

### Refolding Measurements (Equilibrium Refolding Curves)

Equilibrium refolding of γD-crystallin variants was measured using a complementary plate-based protocol in which proteins were first fully denatured at high Gdn with 0.1 mM EDTA and 1 mM DTT and then diluted to lower Gdn concentrations. Denatured protein was diluted into refolding buffer (10 mM phosphate, NaCl at constant ionic strength, 0.1 mM EDTA, 1 mM DTT, pH 7.0) containing 0-5 M Gdn, to a protein concentration higher than 1 µM, and incubated at 37 °C for ∼24 h to ensure complete unfolding.

After this pre-denaturation step, a second 96-well plate (half area UV-Transparent Microplates, Greiner) was prepared with target final Gdn concentrations spanning 0–5 M, using the same buffer composition as in the unfolding experiments. Appropriate volumes of the 5 M Gdn unfolded protein stock were added to each well and diluted with Gdn-containing buffer so that the final protein concentration in every well was 1 µM, the nominal [Gdn] series spanned 0–5 M, and the total volume was 50 µL. Because the unfolded protein stocks contained Gdn, the final denaturant concentration in each refolding well included a constant carryover term proportional to the protein-stock fraction (Δ_[Gdn]_ = [Gdn] _pre-denaturation stock_ × V _pre-denaturation stock_ / 50 µL). Therefore, refolding curves were corrected by shifting the [Gdn] axis to these effective values prior to averaging and fitting. Across three independent runs, the mean carryover corrections were Δ_[Gdn]_ ≈ 0.71 M for NCS, 0.67 M for NCA/T and 0.71 M for WT. Water was added as needed to adjust the final volume while keeping EDTA, DTT, phosphate, and NaCl identical across all wells.

The plate was then sealed and incubated at 37 °C for approximately 24 h to allow the unfolded protein to refold (or remain unfolded) at each Gdn concentration. Intrinsic tryptophan fluorescence was recorded on the same SpectraMax® iD3 multimode microplate reader using excitation at 280 nm and emission scanning from 310 to 420 nm. FI₃₆₀/FI₃₂₀ ratios were calculated for each well as described above and plotted as a function of [Gdn] to generate equilibrium refolding curves. Comparison of unfolding and refolding curves was used to evaluate reversibility and to identify hysteresis and partially refolding intermediates for each γD-crystallin variant.

### Aggregation Assays

Aggregation of γD-crystallin variants was evaluated in a 96-well plate format (half area UV-Transparent Microplates) by monitoring turbidity at 405 nm over time in the presence of various concentrations of guanidine hydrochloride ([Gdn]). Protein samples were prepared at final dimer-equivalent concentrations of NCS 2.5 µM and NCA/T 5 µM in PIPES-based assay buffer (10 mM PIPES, 150 mM NaCl, 0.1 mM EDTA, pH 6.7).

For each condition, degassed protein solutions were dispensed into the wells, followed by addition of a concentrated buffer stock containing EDTA, NaCl, and PIPES to maintain constant ionic strength and chelator concentration across all wells. Gdn stock solutions were then added to achieve final concentrations ranging from 0 to 1.1 M (e.g., 0, 0.1, 0.2, 0.3, 0.4 M, etc.). The remaining volume in each well was adjusted with water to a final volume of 100 µL, ensuring identical total volume and buffer composition across conditions. All reagents, particularly protein and buffer solutions, were degassed prior to use to minimize interference from.

Plates were incubated at 37 °C in a SpectraMax® iD3 multimode microplate reader. Absorbance at 405 nm was recorded every 1 min for a total of 3 h. The time-dependent turbidity traces were used to assess aggregation kinetics at each Gdn concentration, and the maximal slope of each trace was calculated as described (44) and used as the apparent maximum aggregation rate.

### Protein concentration dependence of NCS dimer aggregation at a fixed Gdn concentration

The dependence of aggregation kinetics on protein concentration was examined using covalent NCS dimers in the same 96-well plate format. Aggregation reactions were carried out at 37 °C in the presence of a fixed concentration of guanidine hydrochloride (0.3 M Gdn), under buffer conditions identical to those used in the standard Gdn-dependent aggregation assays (PIPES-based buffer containing NaCl and EDTA).

Purified NCS dimer was diluted into assay buffer to generate a series of stock solutions, which were then dispensed into wells to yield a range of final NCS dimer concentrations (covering the concentrations used in the aggregation experiments shown in **Figure 8E**). Here and throughout, concentrations are reported in terms of covalent NCS dimer; the corresponding monomer-equivalent subunit concentration is twice the dimer concentration. For each condition, protein solution and buffer components were added such that the final composition in every well contained 0.3 M Gdn, constant concentrations of PIPES, NaCl, and EDTA, and a total reaction volume of 100 µL. The volume of water was adjusted as needed to maintain the same final volume across all wells. All protein and buffer solutions were degassed prior to use.

Plates were incubated at 37 °C in a SpectraMax® iD3 multifunctional microplate reader. Turbidity at 405 nm was recorded every 1 min for a total of 3 h. Time-dependent absorbance traces were used to monitor aggregation at each protein concentration. For each condition, the maximal slope of the A₄₀₅ versus time curve was extracted and plotted as a function of NCS dimer concentration to determine the concentration dependence (apparent reaction order) of the aggregation process.

### [Protein]-dependence of NCS dimer aggregation with a fixed [*myo*-inositol]

To examine how protein concentration affects aggregation in the presence of myo-inositol, aggregation assays were performed as described above for the protein-concentration dependence, except that a fixed concentration of myo-inositol was included in all wells. To increase the aggregation rate at low protein concentrations and thereby improve signal-to-noise and kinetic resolution, these assays were performed at 42 °C rather than 37 °C, while all other conditions were kept identical. Purified NCS dimer was diluted into the same PIPES-based assay buffer to generate a series of stock solutions and dispensed into 96-well plates to yield a range of final NCS dimer concentrations (0–5.75 µM) Myo-inositol was added from a concentrated stock to a constant final concentration in every well, while the concentrations of PIPES, NaCl, EDTA and, where applicable, Gdn were kept identical across conditions. The reaction volume in each well was adjusted to 100 µL with assay buffer.

Plates incubated at 42 °C or 37 °C, as indicated, for 3 h in a SpectraMax® iD3 multimode microplate reader. Turbidity at 405 nm was recorded every 1 min. Time-dependent A₄₀₅ traces were used to monitor aggregation at each protein concentration, and the maximal slope of each trace was extracted as the apparent maximum aggregation rate and plotted as a function of NCS dimer concentration in the presence of *myo*-inositol.

### [*Myo*-inositol]-dependence of NCS dimer aggregation at a fixed protein concentration

The dependence of NCS dimer aggregation on myo-inositol concentration was evaluated using the same plate-reader setup, but with a fixed protein concentration and varying myo-inositol. Purified NCS dimer was diluted into assay buffer to a final concentration of 4 µM (dimer basis) in each well. And buffer components (PIPES, NaCl, EDTA) were held constant across all conditions.

*Myo*-inositol was added from a concentrated stock solution to achieve a series of final concentrations (0–0.23 M). The total reaction volume in each well was brought to 100 µL with assay buffer, and all solutions were degassed before use.

Plates were incubated at 37 °C for 3 h in a SpectraMax^®^ iD3 reader, and turbidity at 405 nm was measured every 1 min. Aggregation kinetics were obtained from the OD₄₀₅ versus time curves, and the maximal slopes were plotted as a function of myo-inositol concentration to quantify the effect of myo-inositol on NCS dimer aggregation.

### Transmission Electron Microscopy

Samples for transmission electron microscopy (TEM) were recovered from aggregation reactions and prepared on carbon films on 3- mm 200-nm mesh copper/Pioloform grids (VWR). 10 µl of sample was drop-cast for this purpose. The sample was then stained with 10 μL of 2% (w/v) Phosphotungstic acid (PTA) (Thermo Fisher). Excess liquid was removed, and grids were allowed to dry. The grids were imaged on a FEI BioTwinG2 transmission electron microscope (120 kV).

### Intact Protein Mass Spectrometry

Measurements of protein exact isotopically averaged mass were carried out on a Bruker Impact II QTOF electrospray mass spectrometer equipped with an HPLC system.

## Data Availability

All data are reported within the manuscript.

## Supporting Information

This article contains supporting information: Supplementary Table 1.

## Acknowledgments

The authors are thankful to Drs. David Thorn and Eugene I. Shakhnovich for technical help and advice in the preliminary stages of this project. We also acknowledge Dr. Beniam Berhane and the CASDA Mass Spectrometry Core Facility at Stony Brook University for mass spectrometry experiments.

