## Supplementary material for "Dimerization of natively folded molecules drives misfolding and aggregation in Cys-depleted variants of cataract-associated human lens γD-crystallin": Table S1

**Table S1: Deconvoluted intact mass measurements**

| **NCA/T: 4.98 ml LC peak** | | |  | Expected isotopic average MW: | | | | 20508.7 |  |
| --- | --- | --- | --- | --- | --- | --- | --- | --- | --- |
| **Mass** | **Height** | **Assignment** | **Error, Da.** |  |  |  |  |  |  |
| 20507.8 | **100** | NCA/T native | -0.9 |  |  |  |  |  |  |
| 20553.3 | 9.103 | unassigned |  |  |  |  |  |  |  |
| 20690.9 | 19.897 | contaminant (possibly lipid adduct from LC column) |  |  |  |  |  |  |  |
| 20940.6 | 17.899 | unassigned |  |  |  |  |  |  |  |
| **NCS: 4.98 ml LC peak** | | |  | Expected isotopic average MW: | | | | 20542.6 |  |
| **Mass** | **Height** | **Assignment** | **Error, Da.** |  |  |  |  |  |  |
| 20507.9 | 51.705 | bleed-through from NCA/T sample | -0.8 |  |  |  |  |  |  |
| 20541.9 | **100** | NCS native | -0.7 |  |  |  |  |  |  |
| 20690.9 | 10.103 | contaminant (possibly lipid adduct from LC column) |  |  |  |  |  |  |  |
| 20725.0 | 19.351 | unassigned |  |  |  |  |  |  |  |
| **NCS dimer: 5.01 ml LC peak** | | |  | Expected isotopic average MW (for disulfide dimer): | | | | | 41015.4 |
| **Mass** | **Height** | **Assignment** | **Error, Da.** |  |  |  |  |  |  |
| 20507.4 | 70.35 | bleed-through from NCA/T sample |  |  |  |  |  |  |  |
| 20690.6 | 12.163 | contaminant |  |  |  |  |  |  |  |
| 41014.6 | **100** | NCS disulfide bridged dimer | -0.8 |  |  |  |  |  |  |
| 41055.3 | 20.812 | likely 2x sodium adduct of NCS dimer |  |  |  |  |  |  |  |
| 41102.4 | 17.073 | unassigned |  |  |  |  |  |  |  |
| 41197.7 | 33.04 | contaminant (possibly lipid adduct from LC column) |  |  |  |  |  |  |  |
| 41237.2 | 8.854 | unassigned |  |  |  |  |  |  |  |
| 41285.0 | 6.02 | unassigned |  |  |  |  |  |  |  |
| 41380.4 | 5.278 | unassigned |  |  |  |  |  |  |  |
| **Legend**: Table S1 shows all molecular masses from intact protein LC/MS experiments after automatic deconvolution. All assignments are for isotopically averaged molecular masses. All peak heights are normalized to the highest peak in their respective sample, which is set at 100. All mass assignments are consistent with a calibration error of approx. -0.8 Da. on the mass spectrometry instrument. | | | | | | | |  |  |
